# Natural language signatures of psilocybin microdosing

**DOI:** 10.1101/2022.02.20.481177

**Authors:** Camila Sanz, Federico Cavanna, Stephanie Muller, Laura de la Fuente, Federico Zamberlan, Matías Palmucci, Lucie Janeckova, Martin Kuchar, Facundo Carrillo, Adolfo M. García, Carla Pallavicini, Enzo Tagliazucchi

## Abstract

Serotonergic psychedelics are being studied as novel treatments for mental health disorders and as facilitators of improved well-being, mental function and creativity. Recent studies have found mixed results concerning the effects of low doses of psychedelics (“microdosing”) on these domains. However, microdosing is generally investigated using instruments designed to assess larger doses of psychedelics, which might lack sensitivity and specificity for this purpose. Following a double-blind and placebo-controlled experimental design, we explored natural language as a resource to identify speech produced under the acute effects of psilocybin microdoses, focusing on variables known to be affected by higher doses: verbosity, semantic variability and sentiment scores. Except for semantic variability, these metrics presented significant differences between a typical active microdose of 0.5 g of psilocybin mushrooms and an inactive placebo condition. Moreover, machine learning classifiers trained using these metrics were capable of distinguishing between conditions with high accuracy (AUC≈0.8). Our results constitute first proof that low doses of serotonergic psychedelics can be identified from unconstrained natural speech, with potential for widely applicable, affordable, and ecologically valid monitoring of microdosing schedules.

## 1. Introduction

Psychedelic microdosing consists in consuming relatively small amounts of serotonergic compounds, approximating the perceptual threshold –typically, 10-20% of a full dose (Fadiman & Korb, 2019; Kuypers et al., 2019; Ona & Bouso, 2020; Polito & Stevenson, 2019). Unlike a complete psychedelic dose, microdosing is expected to produce minimal acute effects with sustained effects that can last one or two days. Accordingly, this practice involves interspersing resting days with dosing days (two to four times per week) (Kuypers et al., 2019). Microdosing has been gaining popularity over recent years, with lysergic acid diethylamide (LSD) and psilocybin (in the form of psychoactive mushrooms) being the compounds most frequently consumed for this purpose (Hutten et al., 2019a; Lea, Amada, Jungaberle, Schecke, & Klein, 2020; Polito & Stevenson, 2019; Szigeti et al., 2021). Despite the illegal status of psychedelics in most countries, several websites contain discussions and suggestions related to microdosing and its effects, with users claiming that this practice can improve mood, stimulate productivity, and improve cognitive functions as well as mental concertation (Anderson, Petranker, Rosenbaum, et al., 2019; Lea, Amada, & Jungaberle, 2020). Microdosing is also used for the self-treatment of mental health disorders such as depression or anxiety (Hutten et al., 2019b; Lea, Amada, Jungaberle, Schecke, & Klein, 2020; Lea, Amada, Jungaberle, Schecke, Scherbaum, et al., 2020), and it has been suggested as a model for the clinical use of psychedelics (Kuypers, 2020).

The positive effects of microdosing are supported by multiple observational, survey-based, and open-label studies (Anderson, Petranker, Christopher, et al., 2019; Hutten et al., 2019a; Johnstad, 2018; Lea, Amada, Jungaberle, Schecke, & Klein, 2020; Polito & Stevenson, 2019; Prochazkova et al., 2018; Rootman et al., 2021). However, these generally involve self-selected samples and lack adequate control conditions; thus, given the subtle effects of microdosing, expectations and pre-existing traits may play a fundamental role in the perceived effects (Kaertner et al., 2021; Olson et al., 2020). In contrast, studies following double-blind and placebo-controlled experimental designs have found less support for positive outcomes of microdosing (Bershad et al., 2019; Family et al., 2020; Hutten et al., 2020; Szigeti et al., 2021; van Elk et al., 2021; Yanakieva et al., 2019). Yet, these studies generally include tasks and questionnaires validated in the context of full doses of psychedelics, which might lack the specificity and sensitivity necessary to capture the subtler effects induced by microdosing. In turn, these limitations could hinder the development of translational approaches and the validation of therapeutic models, among other potential limitations, thus raising the need for novel methods to study low doses of psychedelic substances.

We adopted an alternative approach based on natural language processing (NLP). As a first step, this approach consisted in extracting linguistic features from unconstrained speech produced by the participants during the acute effects, using them as input to machine learning algorithms. NLP is characterized for being an objective, non-invasive, cost-effective, and scalable tool to investigate ecologically valid data (Sanz, 2022; Tagliazucchi, 2022). Unlike standard questionaries, which constrain reports to a possibly sub-optimal pre-selected set of questions, this approach is capable of automatically identifying and capturing informative semantic and grammatical features of speech, allowing to distinguish between different experimental conditions (Agurto et al., 2020; Bedi et al., 2014; Corcoran et al., 2018; Norel et al., 2020; Sanz, 2022; Sanz et al., 2021). Importantly, besides informing the contents of the drug-elicited experience, NLP allows to investigate the modulation of language production itself, which can be informative of drug action beyond what is reported during the subjective acute effects (Tagliazucchi, 2022). While NLP has been applied to investigate the effects of different compounds, including psychedelics (Bedi et al., 2014; Cox et al., 2021; Cox & Johnson, 2021; Hase, 2022; Sanz et al., 2021; Sanz et al., 2018; Zamberlan et al., 2018), there are no works to date applying this tool to the study of microdosing.

We analyzed the effects of microdoses of *Psilocybe cubensis* mushrooms (0.5 g dried material, a typical microdose) (Polito & Stevenson, 2019; Szigeti et al., 2021; van Elk et al., 2021) on verbal reports. Data was obtained following a double-blind placebo-controlled experimental design with two different measurement weeks per participant (Cavanna, 2022). During each week, participants received either the active dose or the placebo and were interviewed about their feelings, expectation, perception, mood, creativity, and alertness. We obtained and analyzed two metrics based on our previous results of NLP applied to speech under the effects of high doses of psychedelics: verbosity (total length of the speech samples, in words) and semantic variability (variability of the time series obtained by computing the meaning of consecutive words) (Sanz et al., 2021). We also investigated the mean sentiment score (use of terms linked to positive/negative sentiment) to account for the purported effects microdosing on mood (Cameron et al., 2020; Hutten et al., 2020; Lea, Amada, & Jungaberle, 2020; Lea, Amada, Jungaberle, Schecke, & Klein, 2020; Polito & Stevenson, 2019). Based on these previous reports, we hypothesized that psychedelic microdoses would increase verbosity and mean sentiment scores, and decrease the semantic coherence of speech. Finally, we implemented statistical tests and machine learning tools to classify the experimental condition (active dose of psilocybin vs. placebo) and its unblinding based on these NLP features.

## 2. Methods

### 2.1. Participants

Thirty-four native Spanish speaker volunteers (11 females: 32.09 ± 3.53 years; 23 males: 30.87 ± 4.64 years) were enrolled in this study via social media or word of mouth. All participants enrolled or obtained a higher education degree, had normal or corrected-to-normal vision, and successfully completed all stages of the experiment. Participants reported 11±14.9 past experiences with serotonergic psychedelics of which 1.5±2.3 were considered challenging, and 6 participants reported past experience with microdosing. The following exclusion criteria were assessed after an initial interview to present a general overview of the experiment and sign the informed consent form: past diagnosis of psychotic disorders, bipolar disorders (type 1 or 2), depressive disorders, obsessive-compulsive disorder, anxiety disorder, dysthymia, panic disorder or/and neurological disorders. Those presenting bulimia, anorexia or/and substance abuse/dependence over the last 5 years (excluding nicotine) were also excluded. Participants were asked to stop their habitual use of psychoactive drugs (including alcohol and caffeine) during the duration of the experiment.

This study was conducted in accordance with the Helsinki declaration and approved by the Committee for Research Ethics at the Universidad Abierta Interamericana (Buenos Aires, Argentina), protocol number 0-1054. Speech measurement and analysis was part of a larger protocol pre-registered in www.clinicaltrials.gov, identifier number: NCT05160220. All participants provided their written informed consent to participate in the study. Participants did not receive financial compensation for their participation.

### 2.2. Experimental protocol

This experiment followed a double-blind placebo-controlled design, where two conditions (0.5 g of dried *Psilocybe cubensis* and the same weight of an edible mushroom) were randomized and assigned to two experimental weeks. All participants underwent both experimental sessions. This assignment was done by a third party and blinded to both participants and researchers.

During Wednesday and Friday of each week of the experiment, volunteers attended the laboratory facilities and consumed either capsules with the active dose or the placebo. On Fridays, approximately 2.30 h after dosing, participants were interviewed by a mental health professional member of the research team (FC). The interviews comprised the following questions: “*How do you feel right now?”*; “*Do you feel according to how you expected?”*; “*Are you feeling changes in your perception?”*; “*Are you feeling changes in your mood?”;* “*Are you feeling changes in your level of imagination or creativity?”*; “*Are you feeling changes in your level of attention, alertness or energy?”*. In the following, we refer to these questions as “feeling”, “expectation”, “perception”, “mood”, “creativity”, and “alertness”, respectively. At the end of each Friday, participants were asked to identify the condition corresponding to that week (either active dose or placebo). For a schematic representation of the experimental design, see Figure 1.

**Figure 1.**
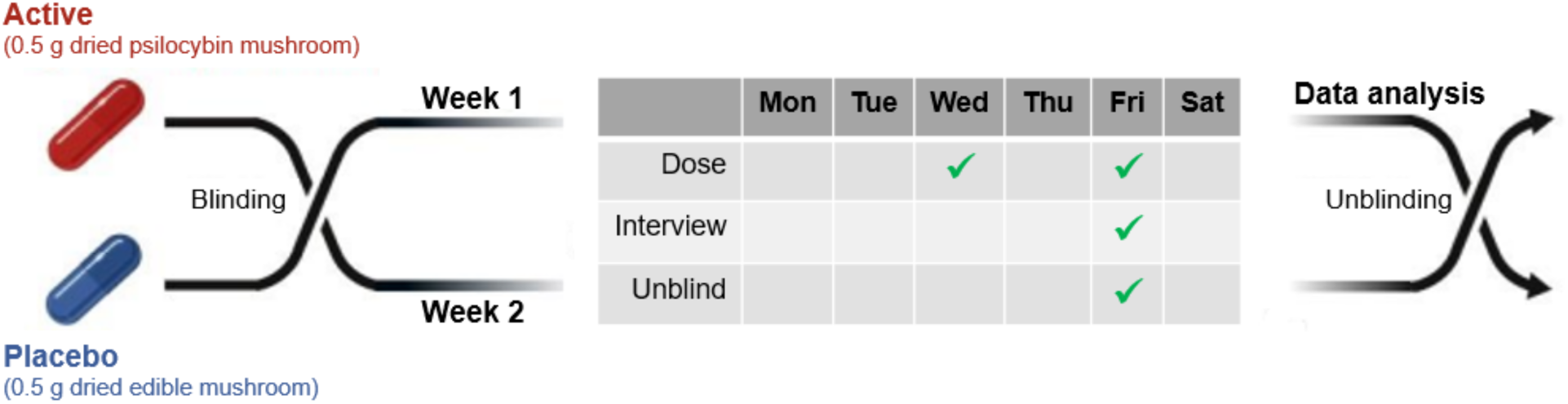
Double-blind placebo-controlled experimental design. The first week, participants were either randomly assigned the active dose or the placebo (in the example shown in the figure, the active condition was assigned to the second week). The remaining experimental condition corresponded to the following week. Subjects and researchers ignored the content of the capsules until the analysis stage. The following schedule was repeated on both weeks: on Wednesday and Friday, participants consumed the dose, and only on Friday subjects were interviewed and asked to guess the condition (unblinding).

### 2.3. Chemical characterization of the mushroom samples

*Psilocybe cubensis* mushrooms from three independent sources were consumed by the participants of this experiment. The active dose was composed of 0.5 g of ground dried material. Samples of 150 mg from each source of material were isolated and sent for analysis (performed by LJ and MK) to the Laboratory of Forensic Analysis of Biologically Active Substances, University of Chemistry and Technology Prague, Czech Republic, which resulted in the concentration of alkaloids psilocybin, psilocin, baeocystin and norbaeocystin, averaged across samples.

Analysis of mushrooms samples revealed the average concentration of the following alkaloids: psilocybin (640.2 μg/g), psilocin (950.7 μg/g), baeocystin (50.4 μg/g) and norbaeocystin (12.5 μg/g). Therefore, the active dose contained 0.32 mg of psilocybin, 0.48 mg of psilocin, 0.025 mg of baeocystin and 0.0063 mg of norbaeocystin. Since at least one month passed between the end of the experiment and the chemical analysis, despite of conserving the samples under optimal conditions, psychoactive material (i.e. psilocybin and psilocin) may have been lost during this time period (Gotvaldova et al., 2021).

### 2.4. Speech analysis

#### 2.4.1. Transcript preprocessing and labeling

Interviews were recorded and manually transcribed by a technician with background in linguistics who was blind to the experimental condition, and afterwards reviewed by a member of the research team as a quality check (both were native Spanish speakers). The parts corresponding to the interviewer were removed from the transcripts, leaving only the parts corresponding to the participants to be analyzed separately for each question.

Each transcript was labeled as active dose or placebo, depending on the experimental condition, and as blinded or unblinded (depending on whether the participant correctly identified the condition). This resulted in four possible combinations (blinded active dose, blinded placebo, unblinded active dose, and unblinded placebo).

#### 2.4.2. Verbosity

We computed verbosity scores by counting the number of words (including repetitions and stopwords) produced by the participants when answering the questions of the interview.

#### 2.4.3. Semantic variability

We computed a metric of semantic variability based on the distance between consecutive words spoken by the subjects. Briefly, this metric was obtained by computing a word embedding for each term in the transcript, with the semantic distance of consecutive words obtained as the cosine between the corresponding vectors in the embedding. Finally, the variability of this time series corresponded to the semantic variability (Sanz et al., 2021). Previous work investigating the acute effects of medium/high doses of serotonergic psychedelics showed that semantic variability is increased during the acute effects relative to the placebo (Sanz et al., 2021; Wießner, 2021). For further details on the implementation of this metric see the supplementary material.

#### 2.4.4. Sentiment analysis

Sentiment analysis is an NLP technique used to determine whether text data conveys positive, negative or neutral sentiment, with applications to quantify the content of experiences elicited by psychedelics (Qiu, 2021). Sentiment analysis of the speech transcripts was implemented in Python using a sentence-level model pre-trained for Spanish (github.com/aylliote/senti-py). Briefly, the model pipeline included standard text pre-processing steps such as conversion to lowercase, accent removal, standardization of linguistic expressions typical of regional Spanish, conversion of verbs to their infinitive forms, introduction of bigrams consisting of adjectives and their negation, and removal of custom words considered useless for the quantification of sentiment. These steps were followed by a univariate feature selector and a Multinomial Naive Bayes classifier trained using labeled data obtained from different websites, comprising approximately 1 million training samples. The parameters and hyper-parameters of this pipeline were found by the use of a grid search combined with K-fold cross validation (K=10). Based on its words, the model assigns to each sentence a score ranging from 0 (negative) to 1 (positive), with a score of ≈0.5 assigned to neutral sentences.

The transcript of each response was first split into sentences, which were used as input for the sentiment analysis model. Finally, the average score across all sentences was computed, resulting in a mean sentiment score (MSS) for the answers to each question in the interview.

The following three sentences extracted from the transcripts illustrate cases with positive, neutral and negative sentiment, according to the model:

*“I feel tired, I’ve been tired all day, I don’t believe that… I keep thinking that I took two placebos”*. Negative sentiment (MSS=0.066).

*“No, I think it was quite normal eh… like overall I feel that I was calmer as when the stuff that normally happens presented itself, I could sort of see it as one step further back you know like more relaxed, but aside from that it was quite normal, I didn’t see any big changes”*. Neutral sentiment (MSS=0.49).

“*Now complete peace of mind”*. Positive sentiment (MSS=0.83).

### 2.4. Statistical analyses

Statistical analyses were conducted using non-parametric Mann-Whitney U tests for the difference in the median of verbosity, semantic variability, and MSS between the groups and subgroups determined by the conditions active dose vs. placebo, and blinded vs. unblinded. The alpha level (0.05) of statistical significance was corrected via the Bonferroni criterion with N=6 (number of questions in the interview). Statistical power analysis estimated values larger than 0.80 for the chosen alpha value and the sample size included in our study (N=34) (Rosner, 2015).

Boxplot representations for each answer and group were used to graphically summarize the results (Figures 2A, 2B, 3A, 3B, S1, S2). Each boxplot comprises from the 25^th^ percentile (lower quartile) to the 75^th^ percentile (upper quartile), whiskers extend from these percentiles up to 1.5 times the interquartile range with a line indicating median values. Participants were represented with single points above the boxplot, except for outliers (below/above the lower/upper quartile ± 1.5 times the inter-quantile range) which were removed.

**Figure 2.**
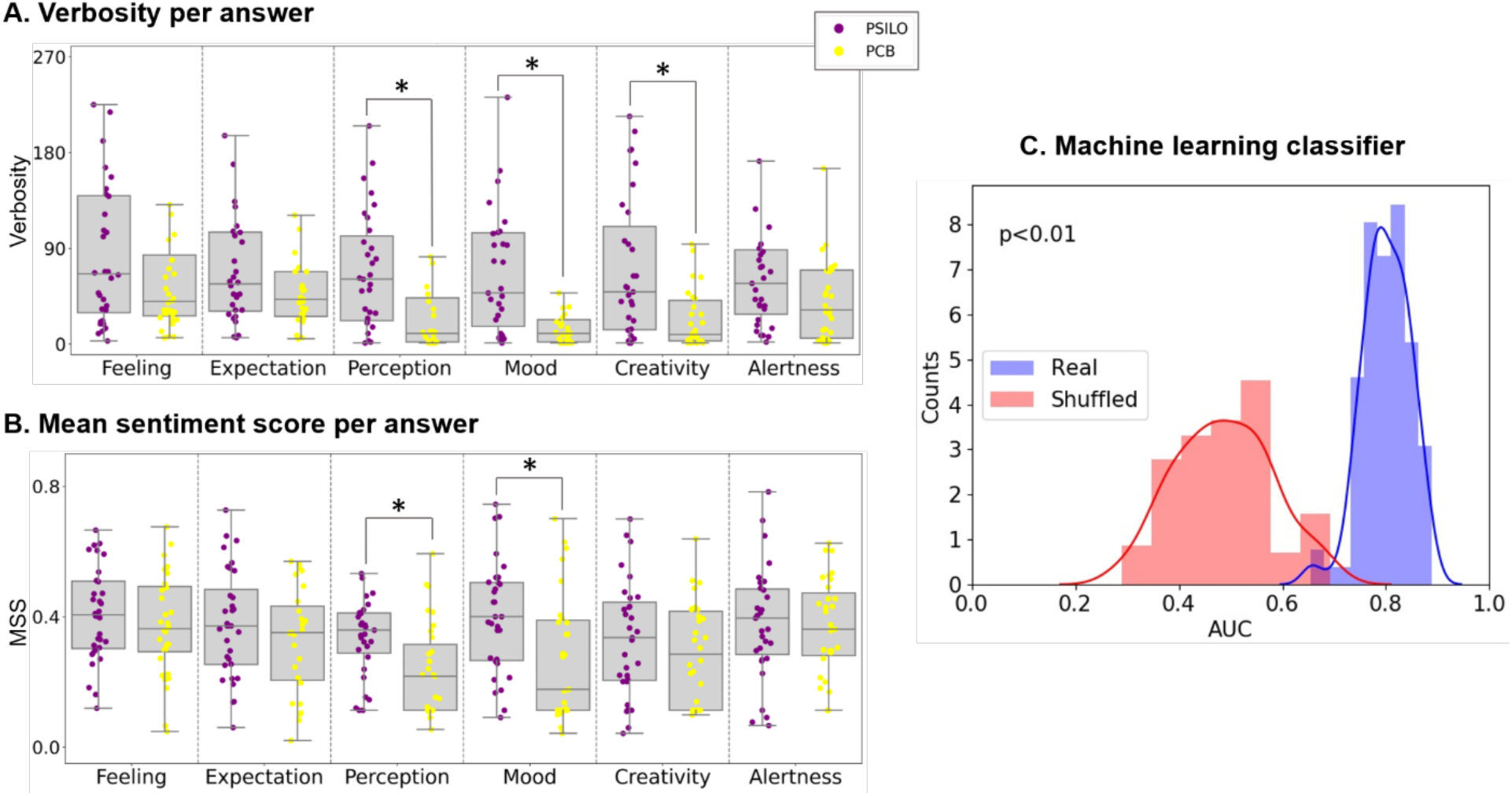
The active dose (PSILO) increased verbosity and positive sentiment relative to the placebo (PCB) **(A)** Boxplots of the verbosity for each question and condition. Significant pairwise differences are indicated with an asterisk (*p<0.05, Bonferroni corrected for multiple comparisons, N=6; Mann-Whitney U test). **(B)** Same information as in panel A but for the MSS. **(C)** Histograms of AUC values obtained from the classification with (red) and without (blue) label shuffling across 100 iterations.

**Figure 3.**
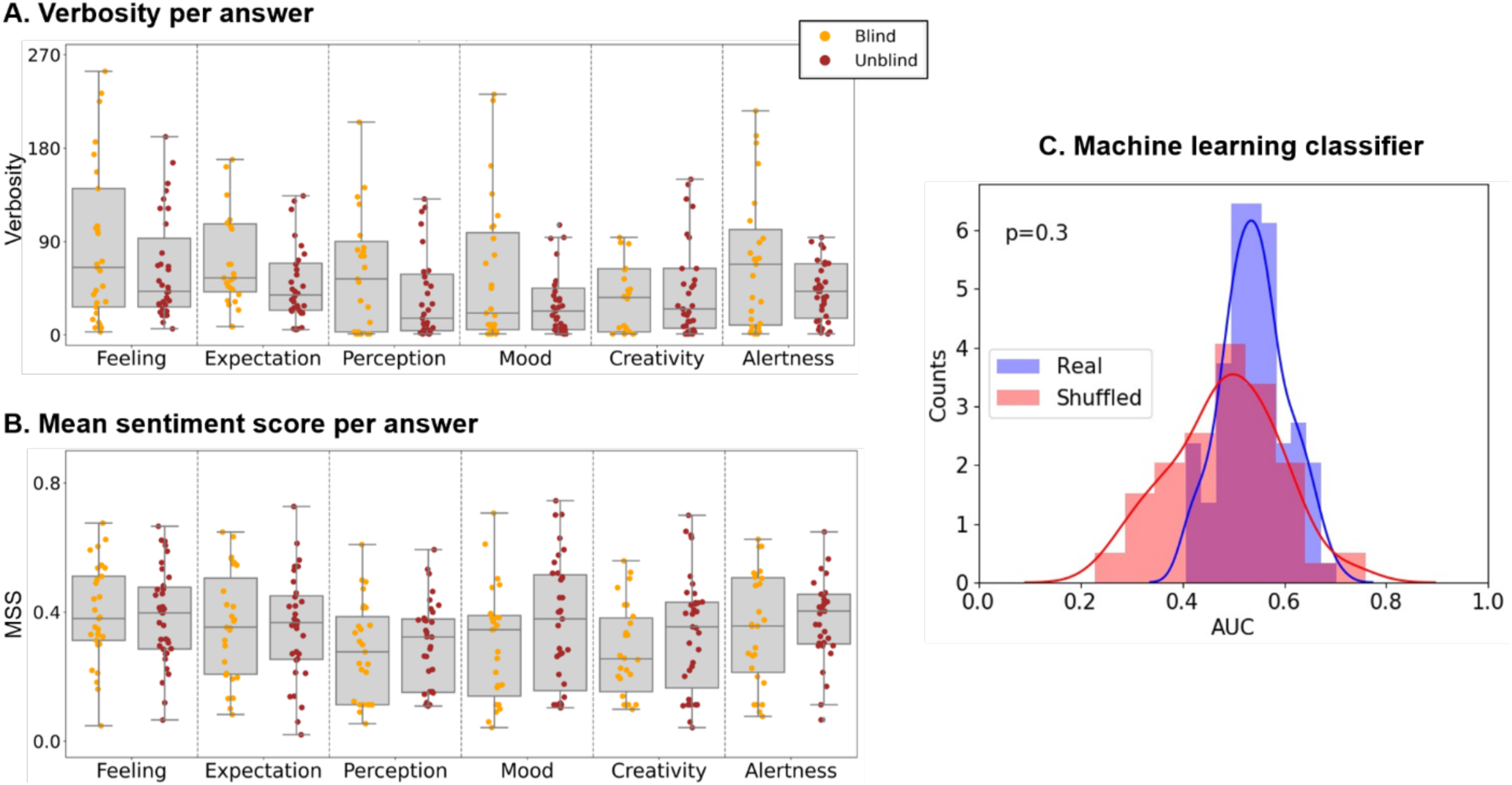
The analysis of verbosity and MSS did not reveal significant differences between blinded (Blind) and unblinded (Unblind) participants. **(A)** Boxplots of the verbosity for each question and condition. Significant pairwise differences are indicated with an asterisk (*p<0.05, Bonferroni corrected for multiple comparisons, N=6; Mann-Whitney U test). **(B)** Same information as in panel A but for the MSS. **(C)** Histograms of AUC values obtained from the classification with (red) and without (blue) label shuffling, across 100 iterations.

Statistical analyses were conducted using Python’s SciPy library (https://scipy.org).

### 2.5. Machine learning analysis

Machine learning models were trained using the scikit-learn (scikit-learn.org) library. In total, each sample consisted of 12 features, given by the verbosity scores and the MSS for each question in the interview (the semantic variability was excluded as it was not significantly different between conditions, see results and supplementary material). Random Forest classifiers (1000 estimators with default parameters, scikit-learn.org) were used to classify these samples according to the experimental condition and its unblinding. This classifier was chosen due to its good performance without need for fine parameter tuning, allowing to avoid the need for a separate testing dataset. In total, six models (same hyperparameters and training-testing procedure) were implemented, two trained to distinguish between experimental conditions (psilocybin vs. placebo) and blinding vs. unblinding, and the remaining four trained to perform the same classification, but restricted to subgroups following the four possible combinations: blinded active dose, blinded placebo, unblinded active dose, and unblinded placebo.

For each classifier, samples were divided into 3 folds via stratified cross-validation to preserve the proportion of labels. Then, each of the folds was used at least once for testing while the remaining were used to train the model; the average of the area under the ROC curve (ROC AUC) across folds was used to determine the model performance. This procedure was repeated 100 times with and without shuffling of the sample labels. This shuffling was used to break the relationship between the features and the labels, thus resulting in a null-model used to estimate chance level performance. Next, the number of times the AUC of the model trained with shuffled labels exceeded the value obtained without label shuffling divided by the total number of iterations (100) resulted in a p-value representing the probability of obtaining the given ROC AUC value assuming chance distribution of the sample labels between classes.

## 3. Results

### 3.1. Statistical analysis and machine learning classification

#### 3.1.1. Active dose vs. placebo

We did not find statistically significant differences in the semantic variability when comparing the psilocybin vs. placebo conditions. For further details see the supplementary material.

Participants under the active dose exhibited higher verbosity than under placebo condition in their answers to all questions in the interview (Figure 2A). Significant differences were found for the answers related to the questions “perception” (U=206, p=0.0012, r=0.41), “mood” (U=177.5, p=0.00022, r=0.47) and “creativity” (U=244, p=0.0077, r=0.32). The remaining answers raised p-values larger than 0.02 and did not survive Bonferroni corrections. For the group under the active dose, median values and interquartile ranges were M=63.5, IQR=82.25 (“perception”); M=50, IQR=85.5 (“mood”) and M=47.5, IQR=92 (“creativity”), and for the group under the placebo condition M=10.5, IQR=39.75 (“perception”); M=10, IQR=19.25 (“mood”); M=11, IQR=40.25 (“creativity”).

Median values of MSS for participants under the active dose were around 0.4, suggesting an overall neutral state (Figure 2B). Participants under the microdosing condition scored higher than under the placebo in all their answers, indicating more positive sentiment. Significant differences were found for answers related to the questions “perception” (U=202.5, p=0.00086, r=0.42) and “mood” (U=243.5, p=0.0076, r=0.33), the other answers resulted in p-values larger than 0.18. Median values and interquartile ranges for the group under the active dose were M=0.37, IQR=0.11 (“perception”); M=0.42, IQR=0.24 (“mood”) and for the group under the placebo condition M=0.22, IQR=0.19 (“perception”); M=0.17, IQR=0.28 (“mood”).

These results suggest that the active dose generated subtle effects leading to longer answers and increased positivity in the speech of the participants when reporting changes to perception, mood and creativity. Answers related to general feelings and expectations remained the same regardless of the experimental condition, as did those corresponding to levels of attention, energy and alertness.

Classification for the instance without label shuffling yielded an AUC value of 0.79 ± 0.04, which was significantly higher (p<0.01) than the one found after label shuffling (0.52 ± 0.10), indicating classifier accuracy above chance level. Histograms representing the AUC values across all iterations are shown in Figure 2C.

#### 3.1.2. Blind vs unblind

Pairwise comparisons between blinded and unblinded participants did not result in significant differences for any of the answers in the interview, yielding p-values larger than 0.013 and 0.1 for the verbosity and MSS, respectively. Participants in the blinded group showed higher median verbosity values (Figure 3A) and lower MSS (Figure 3B) than participants in the unblinded group, but neither of these changes were significant at p=0.05 with Bonferroni correction for multiple comparisons.

The results of the machine learning analysis (Figure 3.C.) were consistent with the lack of statistically significant differences between groups, yielding an AUC value of 0.53 ± 0.06, which did not differ (p=0.37) from the one obtained after label shuffling (0.49 ± 0.11).

#### 3.1.3. Statistical analyses and classifiers restricted to subgroups of participants/conditions

Statistical and machine learning analyses were repeated for all possible combinations of subgroups: blinded active dose, blinded placebo, unblinded active dose, and unblinded placebo. Mann-Whitney U tests were used to compare verbosity, semantic variability and sentiment scores for each subgroup. The performance of the classifiers is summarized in Figure 4. Boxplots corresponding to these results, together with statistical analyses, are included in the supplementary material.

**Figure 4.**
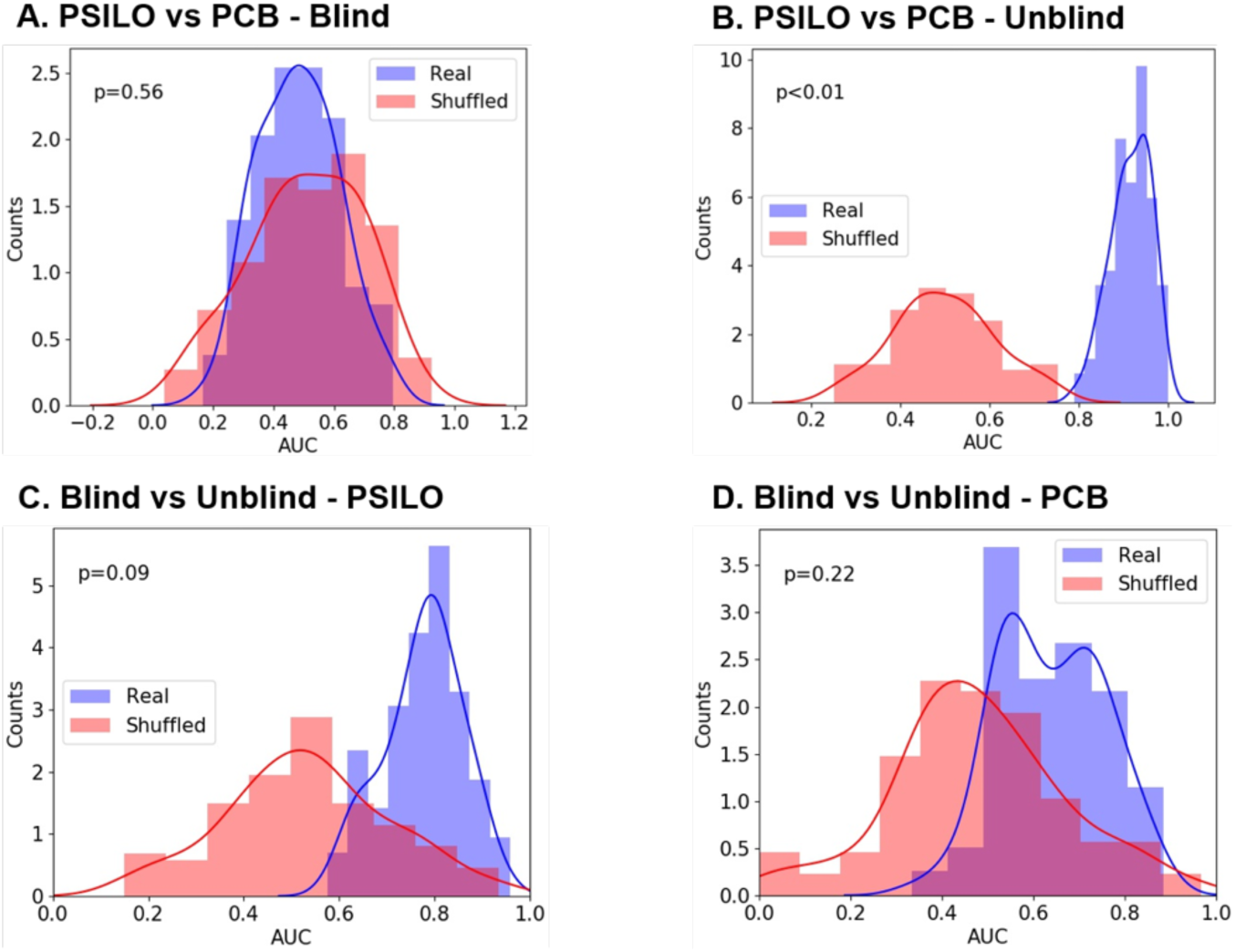
Only the classifier between the active dose and the placebo condition restricted to unblinded participants, yielded a significant accuracy (p<0.05), as compared with the instance with label shuffling. Histograms of the AUC values obtained from classification with (red) and without (blue) label shuffling for all the following subgroups: **(A)** Active dose vs. placebo restricted to blinded participants, **(B)** Active dose vs. placebo restricted to unblinded participants **(C)** Blinded vs. unblinded participants, restricted to the active dose, **(D)** Blinded vs. unblinded participants, restricted to the placebo condition.

The classifier trained to distinguish between the active dose and the placebo condition, but restricted to blinded participants only (Figure 4A), yielded an AUC value of 0.48 ± 0.13. This did not differ significantly (p=0.56) from the value obtained by shuffling labels (0.52 ± 0.18). In contrast, when restricting the samples to unblinded participants (Figure 4B), both conditions were robustly classified as compared to the classifier with label shuffling (p<0.01). AUC values were 0.51 ± 0.17 and 0.92 ± 0.05 for the instances with and without shuffling labels, respectively.

Machine learning results for blinded vs unblinded, but restricted to the active dose and the placebo conditions only (Figures 4C and 4D), did not present statistical differences as compared with the classifiers with label shuffling (p=0.09 and p=0.22, respectively). Nevertheless, classification of blinded vs. unblinded participants limited to the active dose condition presented a p-value close to the alpha level of significance. Under this restriction, AUC values were 0.50 ± 0.18 and 0.76 ± 0.08 for the instances with and without label shuffling. When restricted to the placebo condition, AUC values were 0.50 ± 0.18 and 0.63 ± 0.17, with and without shuffling labels, respectively.

## Discussion

We explored the effects of psychedelic microdosing on natural speech via measures of semantic variability, verbosity, and sentiment scores. The last two measures discriminated between the active dose and the placebo condition. Furthermore, random forest classifiers successfully distinguished between groups based on this information. In contrast, no significant differences in these measures were found between participants who correctly unblinded the experimental condition and those who did not; consistently, machine learning classification yielded an AUC value close to chance level. Finally, statistically significant differences and a robust classification between active dose and placebo were achieved for participants who correctly unblinded the experimental condition.

Previous work has shown that a complete dose of LSD increases verbosity as well as semantic variability (i.e. reduces speech coherence) (Carhart-Harris et al., 2014; Sanz et al., 2021; Wießner, 2021), which is consistent with the general hypothesis of more disordered or entropic brain activity elicited by psychedelics (Carhart-Harris et al., 2014). Insofar as semantic variability indirectly reflects the organization of the stream of thoughts, these previous results suggest that psychedelics result in a hyperassociative state, which in turn might facilitate creativity (Girn et al., 2020). Our results show that semantic variability is not affected by low doses of psilocybin, questioning whether microdosing is capable of enhancing specific aspects of cognition related to creativity through a scrambling effect such as the one postulated for higher doses.

Concerning the results of sentiment analysis, microdosing users generally report an improvement in their mood (Anderson, Petranker, Christopher, et al., 2019; Hutten et al., 2019a; Johnstad, 2018; Lea, Amada, Jungaberle, Schecke, & Klein, 2020; Polito & Stevenson, 2019; Prochazkova et al., 2018; Rootman et al., 2021). Accordingly, increased MSS of natural speech could reflect the positive effect of psilocybin on mood and subjective well-being. Interestingly, increased MSS was not only observed in the answer to the question about mood included in the interview, but also in the answer to other questions. This suggests that microdosing could be capable of inducing a state of positive mood, which generally affects verbal expression, and might be indicative of improved mental health (Babu & Kanaga, 2022). The same applies to increased verbosity, which could reflect more enthusiasm, motivation and energy during the acute effects of the microdose (Sanz et al., 2021). Considered separately, these results are compatible with alternative scenarios such as increased verbosity due to nervousness, or increased MSS due to the inclusion of positive terms in the speech that are not directly implicated with the mood of the participants. However, when considered together they support a synergetic interpretation favoring the induction of a state of positive mood, even though additional experiments should be conducted to exclude alternative interpretations.

The comparison between blinded and unblinded participants suggests that expectations play an important role in the perceived effects of microdosing. This could explain the lack of significant differences and poor classifier performance in the comparison between placebo and active dose, restricted to participants who did not identify the experimental condition. However, expectation effects were not apparent in the comparison between blinded and unblinded participants restricted to the active dose, suggesting that microdosing could generate effects that cannot be fully explained by the identification of the experimental condition. The issue of unblinding is pervasive to all studies of psychedelic microdosing and, more generally, to the study of compounds capable of eliciting profound alterations in the state of consciousness (Kuypers et al., 2019; Muthukumaraswamy et al., 2021; Schenberg, 2021; Szigeti et al., 2021; van Elk et al., 2021). Future studies should explore more adequate control conditions and experimental paradigms capable of alleviating these concerns.

Natural language processing tools allowed us to reveal significant differences and to obtain a robust classification of the conditions without resorting to questionaries that were not formulated with the specific objective of studying low doses of psychedelics. Other advantages of these methods include scalability, low implementation cost, and capacity to process large volumes of unstructured data produced under conditions of ecological validity (Tagliazucchi, 2022). Because of these advantages, NLP tools could be useful to provide long distance guidance for individuals following future therapeutic protocols with serotonergic psychedelics (Kuypers, 2020). On the other hand, the process of conducting interviews could be lengthy and time consuming, representing a limitation unless novel tools to automate the process are developed.

Our study presents some limitations stemming from its experimental design. We considered microdosing effects over periods of one week; thus, long-term outcomes associated with cumulative effects cannot be captured by our speech analysis. However, since NLP measures can be extracted remotely and automatically, this issue could be mitigated by asking participants to self-record short speech samples and then to submit them for analysis. This observation also highlights the potential applicability of these findings as a tool to monitor the effects of microdosing based on samples of ecological validity. While we only considered semantic variability, verbosity (which are increased by higher doses of psychedelics) (Sanz et al., 2021) and the MSS (due to reports of mood enhancements induced by microdosing), future studies should explore other specific measures that could more adequately capture the effects of microdosing on cognition and mental function. Also, the acoustic analysis of speech samples (e.g. prosody) could yield valuable information, but remains largely unexplored in the context of psychedelics, both for low and high doses (Tagliazucchi, 2022).

In conclusion, we characterized natural language produced under effects of low doses of psilocybin, extracting markers from unstructured and unconstrained speech that are compatible with improved mood of the participants, and which might be difficult to capture using more traditional methods. These results highlight the value of recording brief samples of natural language before, during and after the acute effects of psychedelic compounds.

## Acknowledgments

We thank Sol Pérez Vázquez for her assistance with the logistics of this study. This study was funded by grant PICT-2019-02294 awarded by Agencia Nacional de Promoción Científica y Tecnológica (Argentina). Partial funding was received from the Ministry of Health of the Czech Republic (grant no. NU21-04-00307) and the Grant Agency of the Czech Republic (grant no. 20-25349S). Adolfo García is an Atlantic Fellow at the Global Brain Health Institute (GBHI) and is supported with funding from GBHI, Alzheimer’s Association, and Alzheimer’s Society (Alzheimer’s Association GBHI ALZ UK-22-865742); ANID, FONDECYT Regular [1210176]; and Programa Interdisciplinario de Investigación Experimental en Comunicación y Cognición (PIIECC), Facultad de Humanidades, USACH.

