## Supplementary Material for "Natural language signatures of psilocybin microdosing"

*Analysis of subgroups (placebo vs. active dose, blinded vs. unblinded)*

Boxplot representations of the verbosity and MSS for all possible combinations of placebo/active dose and blinded/unblinded are shown in Figure S1. Participants under the active dose scored higher median values of verbosity and MSS than under the placebo, regardless of being blind or unblind to the experimental condition. However, significant differences were found only in the unblinded group since all answers of the blinded group yielded p-values larger than 0.066 and 0.022 for the verbosity and MSS, respectively. For the verbosity measure, significant differences were found for questions related to “perception” (U=27.5, p=1.7x10$^{-5}$, r=0.7), “mood” (U=36.5, p=6.3x10$^{-5}$, r=0.65), “creativity” (U=58, p=0.0009, r=0.53) and “alertness” (U=72.5, p=0.004, r=0.45); the other answers raised p-values larger than 0.022. Median values and interquartile ranges for the group under the active dose were M=58.5, IQR=90.8 (“perception”); M=45, IQR=66.8 (“mood”), M=51, IQR=94.5 (“creativity”) and M=56.5, IQR=50 (“alertness”). The group under the placebo condition scored M=5, IQR=9 (“perception”); M=5, IQR=15 (“mood”), M=9, IQR=27 (“creativity”) and M=28, IQR=40 (“alertness”). For the MSS measure, significant differences were found in answers related to “perception” (U=52.5, p=0.0005, r=0.56) and “mood” (U=75, p=0.005, r=0.44), the other answers yielded p-values larger than 0.18. Median values and interquartile ranges for the group under the active dose were: M=0.38, IQR=0.09 (“perception”) and M=0.44, IQR=0.15 (“mood”), while the group under the placebo condition scored M=0.22, IQR=0.17 (“perception”) and M=0.14, IQR=0.29 (“mood”).

Comparisons of blinded vs. unblinded participants restricted to either active dose or placebo did not result in significant differences. Restriction to the active dose condition yielded p-values larger than 0.023

(verbosity) and 0.022 (MSS). Under the PCB condition p-values were larger than 0.023 (verbosity) and 0.085 (MSS).

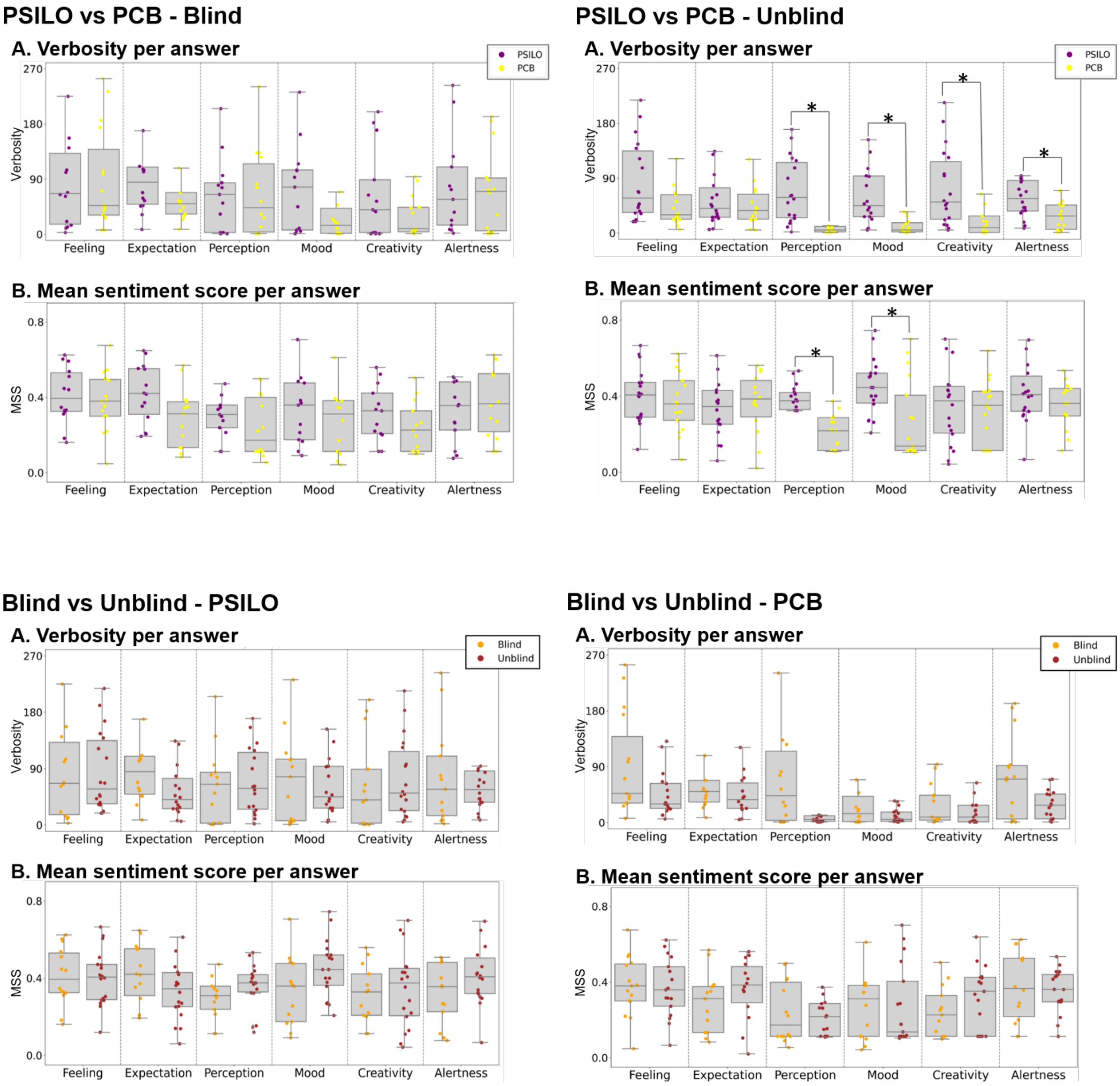

**Figure S1.** Only the comparison between active dose (PSILO) vs. placebo (PCB) restricted to unblinded participants resulted in significant differences. Boxplot representations of the verbosity **(A)** and MSS **(B)** measures for active dose vs. placebo restricted to blinded participants (top left) and to unblinded participants (top right); blinded vs. unblinded participants restricted to the active dose condition (bottom left) and to the placebo condition (bottom right) (*$p<0.05$, Bonferroni corrected for multiple comparisons, N=6; Mann-Whitney U test)

*Semantic variability*

To compute the semantic variability, we first applied a series of preprocessing steps including splitting the document into its individual words, removal of punctuation marks and other symbols, conversion to lower case, removal of stopwords, lemmatization (i.e. conversion of each word to its base form). Next, we used a word embedding method (FastText, https://fasttext.cc) to obtain a vector representation of each word, allowing to compute semantic distance between words as the cosine between the corresponding vectors (Corcoran et al., 2018). From this embedding, we computed the semantic distance between consecutive words and then obtained the variance of this sequence of numbers as a metric of semantic variability (frequent jumps in semantic distance imply high variance and vice versa) (Sanz, 2022; Sanz et al., 2021). Results of this analysis are shown in Figure S2.

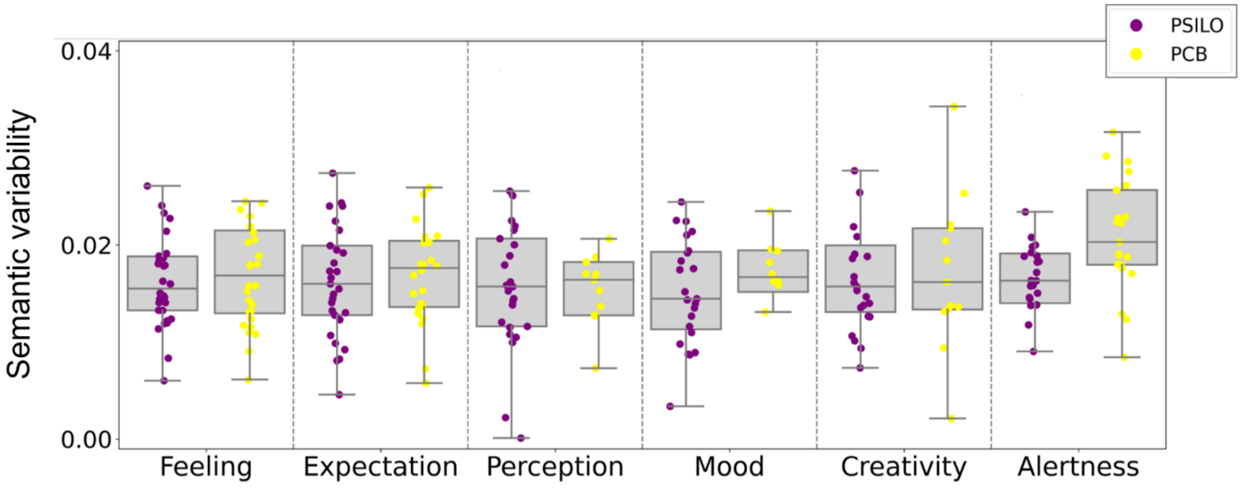


**Figure S2.** Semantic variability did not result in significant differences between active dose (PSILO) and placebo (PCB), all p-values were larger than 0.0092. Boxplot representations of this metric for each question and condition.
